# Stability metrics behave predictably across data qualities but are sensitive to community size

**DOI:** 10.1101/2023.09.25.559298

**Authors:** Duncan A O’Brien, Christopher F Clements

## Abstract

Modern biodiversity monitoring is generating increasingly multidimensional representations of wildlife populations and ecosystems. It is therefore appealing for conservation and environmental governance to combine that information into single measure of ecosystem or population health. Stability represents a desirable feature of ecosystems that supports this aim, measured through resistance, recovery, and variability. In deterministic mathematical systems, the Jacobian matrix is a common characteristic used to quantify resistance and resilience and whilst historically it has been challenging to estimate from empirical data, recent work has proposed a suite of metrics capable of reconstructing it for a real-world community using time series data. Here we assess the robustness of three Jacobian metrics and two variability estimating stability metrics to varying time series lengths and data qualities based on that seen in real-world wildlife time series. Using Lotka–Volterra equations, we generate short time series (to match global biodiversity datasets such as the Living Planet Index and BIOTIME) and introduce sampling error corruptions (to mimic varying search efforts) to validate metric performance in empirical data. The robustness stability metrics generally improved with time series length and search effort in the anticipated manner. However, number of species dramatically altered metric capability, with larger communities decreasing the reliability of stability metric trends. Overall, stability metrics behave predictably across realistic data corruptions. Generic stability estimation is therefore possible from abundance time series alone, and we suggest that, given the increasing availability of multivariate community data, focussing on Jacobian estimates is a plausible ecosystem condition indicator.

## 1. Introduction

Biodiversity loss is a major concern for the maintenance of ecosystem stability and functioning (Cardinale et al., 2012; Oliver et al., 2015). Despite this, species extinctions are accelerating (Ceballos et al., 2015) with repercussions for ecosystem services, exploitable resources, and habitat cover (Traill et al., 2011; Tylianakis et al., 2008). In combination, these negative impacts have resulted in biodiversity policy targets being commonplace and statutory (United Nations, 2020). The conservation and political decisions behind the selection of these targets is dependent on sufficiently high quality data and reliable indicators of biodiversity change (Dale and Beyeler, 2001). Such biodiversity indicators are ultimately used in an ‘indicator-policy cycle’ (Nicholson et al., 2012) which quantifies the scale of impacts and monitors whether the desired outcomes have been achieved (Donatti et al., 2020). This is critical to ensure appropriate policy decisions during a period of competing priorities for decision makers.

The Living Planet Index (LPI - Loh et al., 2005) has emerged alongside BIOTIME (Dornelas et al., 2018), the Global Population Dynamics Database (Inchausti and Halley, 2001) and the IUCN Red list (IUCN, 2022) as the focal datasets and descriptive instruments for global biodiversity trends. They have influenced international (United Nations, 2020, 1992), national (Environment and Climate Change Canada, 2019), and regional (Brotons et al., 2020) environment policy, as well as scientific discourse (Dornelas et al., 2019; Gonzalez et al., 2016; Johnson et al., 2024; Leung et al., 2020; Loreau et al., 2022; McRae et al., 2017). It is therefore undeniable the importance of these datasets for current and future biodiversity monitoring especially given the relative paucity of long-term and reliable counts of species globally (Hughes et al., 2017; McRae et al., 2017). Consequently, the available time series have driven a primary focus on abundance changes by biodiversity indicators in absolute or relative terms (i.e. species richness or diversity indices), despite the acknowledged sampling errors, geographical biases, and many short time series present (Gonzalez et al., 2016; Leung et al., 2020). These concerns generate uncertain estimates of global biodiversity change (Johnson et al., 2024; Wauchope et al., 2019) which hamper the indicator-policy cycle.

Such a focus on abundance also potentially overlooks other signals expressed by stressed populations which are likely mechanistically important for abundance decline and can provide earlier and more robust indications of biodiversity loss. Unfortunately, for many practitioners, abundance data is all that is available and limits which indicators can be calculated. One suggestion is that ecosystem complexity and stability are more informative than raw abundance trends (Capdevila et al., 2022; Pimm, 1984). Here we define complexity and stability following Pimm (1984) as ‘the composition of a system’, and the capacity to ‘return to the [system’s] initial equilibrium [or state] following [it] being perturbed from it’ respectively. Complexity is routinely represented by species richness and evenness while measuring stability is challenging (Capdevila et al., 2022; Donohue et al., 2016, 2013) with an ever expanding suite of possible metrics and indices proposed (Kéfi et al., 2019). Despite this, stability is often more informative than complexity for a practitioner as most managed systems are at an equilibrium/stable state that provides a beneficial function – be that an ecosystem service or another property – and any deviation from stability risks compromising that function. Further, the evidence around complexity responses to stress is unclear (Pimm, 1984), whereas stability is anticipated to decline. Consequently, empirically estimating the response of the ecosystem to stress as a measure of population/ecosystem health rather than solely its species composition has merit.

Stability is by definition (Pimm, 1984) multidimensional (Figure 1) with three primary axes – variability, resilience and resistance. Kéfi et al. (2019) and Donohue et al. (2013) specifically highlight that these dimensions are rarely considered in parallel, across neither population nor community scales. Empirical stability research should therefore attempt to reconcile such differences in scale and dimensionality. In addition, Kéfi et al. (2019) also suggest that any metric must be rooted in theory and be consistent across empirical and theoretical studies. For example, the dominant eigenvalue of the Jacobian is common in theoretical studies as a measure of resilience/asymptotic stability (i.e. the ability of a system to return to a steady state)(Baruah et al., 2022; Dakos, 2018; Strogatz, 2015), but has rarely been reported in real-world studies (Kéfi et al., 2019) due to the challenge of defining empirical Jacobians.

**Figure 1.**
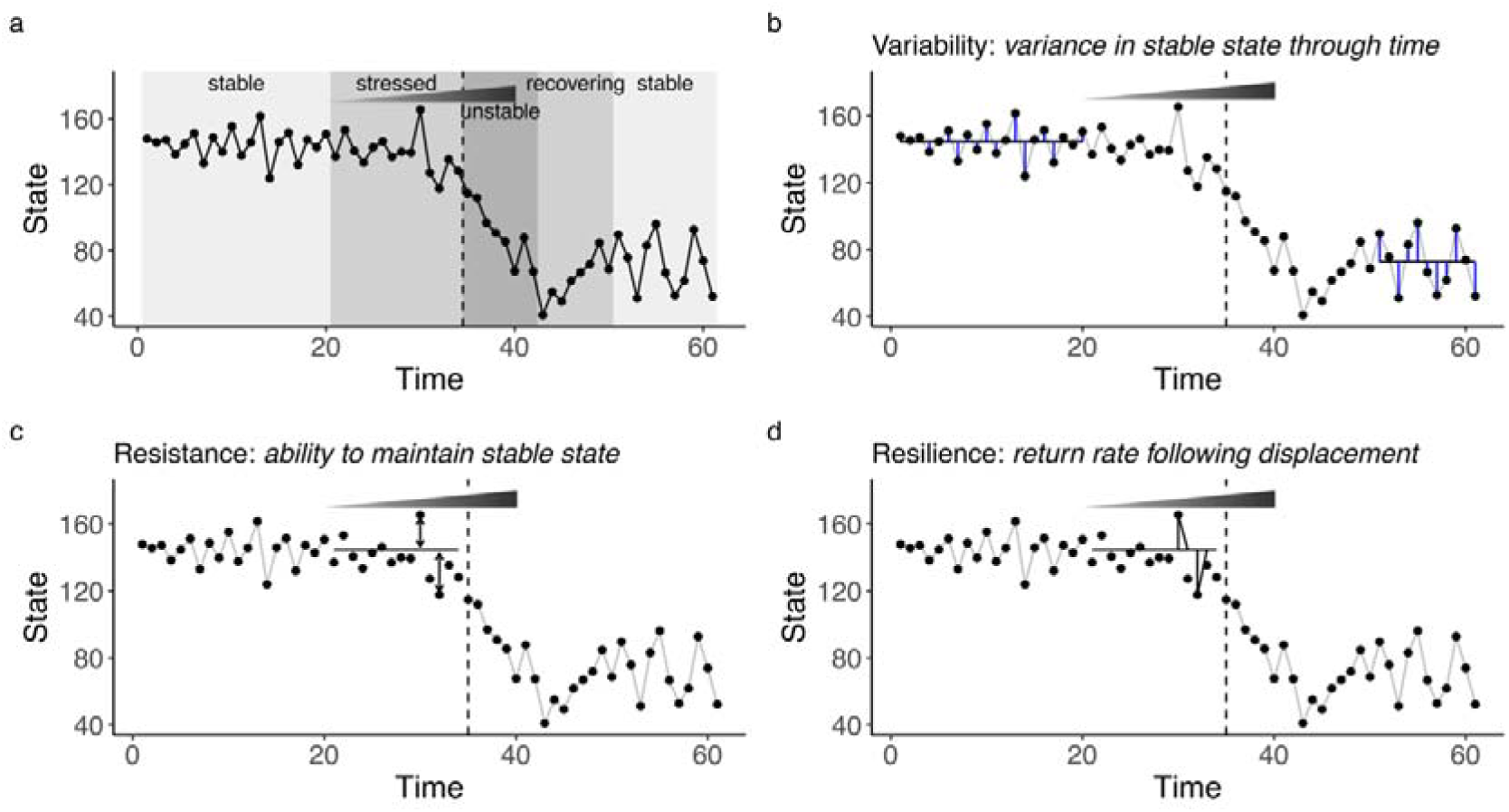
The three primary dimensions of stability defined by Pimm (1984) and how they relate to a system experiencing a ramp stress. a) An example system where state is changing through time in response to a ramp stress. Points and lines represent a unidimensional characteristic of this system (total biomass for example), and the shaded triangle indicates the period of time where a stress is being gradually increased. The system begins in one stable state, experiences the stress, undergoes a rapid transition (a.k.a a tipping point) after which the system stabilises in a new stable state. These discrete periods are indicated by shaded bars and the tipping point is marked with a dotted vertical line. b) The variability dimension of stability represents the inherent fluctuation(s) (blue lines) around the stable state(s) (solid horizontal lines). c) Resistance is how far from the stable state average (solid horizontal line) does the stress value push the system. d) Resilience represents how quickly (the angle of the triangles) does the system return to its stable state average (solid horizontal line).

Computational advancements in the last decade have provided a novel toolbox for estimating Jacobians and species interaction matrices from observational data (Ahmad et al., 2016; Sugihara et al., 2012; Williamson and Lenton, 2015). These indicators are grounded in dynamical system theory (Strogatz, 2015) and have the potential to quantify the resilience dimension of stability under strong equilibrium but also under more complex and chaotic dynamics (Lyapunov, 1992). This is obviously attractive for biodiversity monitoring as loss of resilience/stability is indicative of oncoming decline (Dakos et al., 2015) even if the system is temporarily away from equilibrium (which is likely the case for ecosystems experiencing multiple stressors (Hastings et al., 2018)). However, it is unclear how stability metrics behave across data qualities and ecosystem community structure to warrant their immediate usage in real-world data. Similar ambiguities have influenced the practicality of related population collapse indicators (a.k.a. Early Warning Signals – EWSs) with good performance reported in idealised simulated studies (Clements et al., 2015; Dakos, 2018) but failures in observational time series (Burthe et al., 2016; O’Brien et al., 2023a). There is consequently a need to test the behaviour of the stability metric toolbox in controlled scenarios matching the data qualities present in biodiversity datasets such as the LPI, and ensure their behaviour is sufficiently reliable prior to their potential usage in the indicator-policy cycle.

In this work, we use simulated multispecies ecological communities to test the influence of time series length and data corruption upon five stability metrics in a system where the true dynamics are known. These stability metrics included a multivariate index of variability (a measure of variability - (Brock and Carpenter, 2006)), Fisher information (a measure of variability and resistance -(Ahmad et al., 2016; Karunanithi et al., 2008)), and three Jacobian matrix estimators (measures of resilience - (Grziwotz et al., 2023; Ushio et al., 2018; Williamson and Lenton, 2015)). We specifically estimated the trend of each metric in 5, 15 and 25 species communities across different scales (populations vs communities) and show that while metric reliability generally increases with data quality, metrics displayed weak but correct trends even in low data qualities. However, the choice of species contributing to the metric dramatically influenced reliability leading to recommendations that users must guide their species selection with ecological knowledge.

## 2. Methods

### 2.1 Simulation framework

Community dynamics were simulated using an established multi-species generalised Lotka-Volterra competitive model (Hofbauer and Sigmund, 1998), containing a control parameter that can drive one or more species to extinction. Specifically, this model simulates a species beginning to outcompete others (Equation 1 - (Dakos, 2018)) in an established competitive community and is written as:

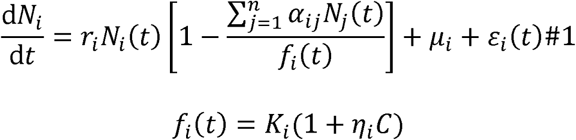

where *i* and *j* are the focal and interacting species identities respectively. *N_i_* is the resulting population density of species *i*, *r_i_* is *i*’s reproduction rate, *α_ij_* is the interaction coefficient between species *i* and *j* (or the per capita effect of species *j* on *i*), *f_i_* is the species-specific out-competition function with *C* the control parameter (i.e. the application of stress to the community), and *ε* is a white noise process with mean 0 and variance 0.1. Symbols *K_i_*, *μ*_i_, and *η_i_* represent the initial carrying capacity, immigration term, and species-specific response to the control parameter *C* for species *i* respectively. Altering *C* directly influences a species’ carrying capacity *K* which alters its ability to compete.

The community parameters were parameterised followed Dakos (2018). Specifically, all interaction matrix *α_ij_* elements were considered competitive and drawn from a uniform distribution 0 – 1.5. The diagonal was set to 1 to ensure strong intraspecific competition. Reproduction rate *r*_i_, carrying capacity *K*_i_, and sensitivity to the control parameter *η_i_* were sampled from the uniform distributions 0.9 – 1.1, 5 – 15, and 0 – 1 respectively. The immigration rate *μ*_i_ was set as 0.01 for all species. We consequently generated 25 communities with unique interaction matrices.

Each community was stochastically simulated using the Euler-Maruyama method at d*t* = 1, for 100 timesteps with a control parameter value of 0. In the following period between t100 and t200, the control parameter value was incrementally increased for one species and then held at the maximum value of 5.0 (Dakos, 2018) as a ramp stress from t200 to t300. This results in the carrying capacity of the selected species being raised and that species’ ability to out-compete its rivals increasing to a threshold where the entire community becomes unstable.

The out-competing species within each community was chosen at random to generate a diversity of community dynamics. Each community was simulated 25 times repeated across stressed and unstressed regimes resulting in 1250 timeseries per community size (2 stress regimes, 25 communities, 25 stochastic runs). Simulations were run using the ‘*DifferentialEquations*’ package (Rackauckas & Nie 2017) in the Julia language (Bezanson *et al*. 2017).

To clarify, the purpose of these simulations is to create a library of stressed community dynamics where the true fate is known for indicator testing. The choice of this simulation framework ensures that the shape and trends of the community dynamics can be both linear and/or non-linear (Dakos, 2018) akin to observed wildlife time series (Johnson et al., 2024; Ledger et al., 2023), and so are appropriate to challenge these proposed stability indicators.

### 2.2 Data post-processing

Each simulated time series was then altered to mimic the imperfect sampling and short time series found in the LPI and BIOTIME. First, we identified the time point of fastest change in the community’s first principal component using the bisection method (Christopoulos, 2016) provided by the *inflection* R package (v.1.3.6). This method simply estimates the first derivative between successive points and searches for the largest derivative. Identifying these changes ensures that the community is stressed but no species has been yet driven to collapse (Figure 1b), and allows us to identify practical stability metric behaviour over timescales appropriate for intervention (Davidson et al., 2018; Hastings, 2016). Consequently, each time series was truncated to 10, 25, 40, 55 and 70 timepoints prior to the estimated inflection point, matching the range of time series lengths contributing to the LPI/BIOTIME. Each truncated time series was then subjected to simulated searches ranging from 10 to 100% effort in 10% increments. To achieve this, we converted the simulated continuous species biomass data to counts, to represent natural animal populations, by multiplying the simulated data by 100 and rounding to the nearest integer. We then drew ‘observed’ species counts from a binomial distribution with probability equal to the search effort (Clements et al., 2015), which mimics both imperfect spatial sampling and detection. Resultantly, each initial simulation was reanalysed for each time series length and search effort combination. All post processing, metric calculation and analysis was performed in the R language v.4.3.0 (R Core Team, 2022).

### 2.3 Stability metrics

Five stability metrics were calculated for each post-processed simulation using the R package EWSmethods (v.1.2.0) (O’Brien et al., 2023b). These metrics included Fisher information (Karunanithi et al., 2008), a multivariate index of variability (Brock and Carpenter, 2006), two S-map estimated Jacobian indices (Grziwotz et al., 2023; Ushio et al., 2018) and a Jacobian index estimated from multivariate autoregressive models (Williamson and Lenton, 2015). All metrics were calculated in a rolling window 50% the total time series length following the typical practice for generic indicators (Dakos et al., 2012; Lenton et al., 2012). A visualisation of each metric’s calculation procedure and anticipated behaviour under different data qualities is available in Figure S1.

Fisher information (FI) quantifies ‘indeterminacy’ or ‘the amount of information observed data contain on the unmeasured state’ (Ahmad et al., 2016). It is estimated as the probability a the community is in the same state as the previous time point. Typically, a ‘state’ is defined by the variance of each time series in a reference period (Karunanithi et al., 2008). We therefore estimate FI in a rolling window along the time series, with the variability of each window compared to that of the previous. If the variability is greater than that of the reference variability, then the windows are binned into separate states. Here we defined this reference variability as the standard deviation of each species across the entire time series (Karunanithi et al., 2008). The FI value is then the amplitude or height of the probability distribution that a window may be in any one of the possible states. Fisher information consequently is a representation of variability and resistance (Figure 1b,c) and is expected to decrease along with stability.

The multivariate index of variability (mvi) is the square root of the species covariance matrix’s dominant real eigenvalue and represents a multivariate equivalent to the univariate “variance” early warning signal (Brock and Carpenter, 2006). This eigenvalue represents multivariate variance analogously to a principal component– where the associated eigenvector of the dominant eigenvalue is the line that explains most of the variance in the multivariate dataset. The multivariate index of variability is a representation of variability (Figure 1b) and is expected to increase as stability decreases.

The two S-map Jacobian indices are derived from a state space reconstructed via empirical dynamic modelling (Park et al., 2021; Sugihara et al., 2012). In brief, both indices estimate the Jacobian of the community by comparing the time delayed relationships between each representative time series and predicting out from the reconstructed state-space (Ushio et al., 2018). The coefficients of the local linear model performing this out-prediction represent a time-varying estimate of the community’s interaction matrix (*α_ij_*), from which the real part of the dominant eigenvalue is the community’s “asymptotic stability”. These metrics quantify the resilience dimension of stability (Figure 1d). All species data were scaled to mean zero and unit variance prior to metric estimation, following Chang et al (2017), to ensure all species are of equivalent magnitude to maximise the accuracy of the S-map estimated interaction strengths.

The multivariate form of this Jacobian index (multiJI) compares all pair wise interactions between species in the community and constructs the state space by delay embedding up to *E*, where *E* is the number of species. The metric is therefore sensitive to the choice and number of variables (i.e species here) contributing to the S-map calculation (Medeiros and Saavedra, 2023; Ushio et al., 2018), and limits the minimum number of time points to *E*. In this study, as all species are contributing to system stability, all species are appropriate for inclusion.

However, if a species count is constant for the duration of a window, then S-map reconstruction is impossible and therefore we excluded any species which were unchanging for the duration of the time series. This is possible in our simulations when a species either goes extinct prior to the shortened time series assessed, or low search efforts did not encounter species with small populations. Similarly, as time series length was often less than *E* in the specious communities (e.g. 10 timepoints but *E* = 15), for these communities we randomly selected species to contribute to multiJI to test this additional processing error on the capability of multiJI to accurately characterise stability change. multiJI is expected to increase as stability decreases with a value greater than 1 indicating instability, whereas a value below 1 indicates stability.

The univariate form of Jacobian index is calculated for each species independently by delay embedding the time series against lagged versions of itself (Grziwotz et al., 2023). The resulting local linear model coefficients (i.e *α_ij_*) are therefore all 0, excluding the lower-off diagonal (all −1), whereas the first row elements become the coefficients for the focal species. Consequently, the system’s behaviour can be approximated in the absence of complete information and escapes concerns regarding the inclusion of variables/species (Grziwotz et al., 2023; Medeiros et al., 2022; Ushio et al., 2018). Here we set the embedding dimension *E* to 1 (1 “year”) and the time lag *τ* to 10% the length of the time series following the coded examples of Grziwotz *et al*. (2023). To allow comparability with the multivariate stability metrics, we then averaged uniJI estimates across all species (mean_uniJI) and extracted the maximum index value observed (max_uniJI), with both metrics also expected to increase in parallel with stressor value similar to mvi and multiJI.

Finally, the multivariate autoregressive Jacobian index (multiAR) fits autoregressive models between multiple time series and uses the generated autocorrelation matrix’s eigenvalues to estimate the system’s Jacobian dominant eigenvalue (Williamson and Lenton, 2015). The eigenvalues of the two matrices’ real components are related by the equation:

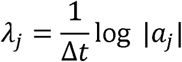

where λ*_j_* is the Jacobian eigenvalues, *a*_j_ is the eigenvalues of the autocorrelation matrix and Δ*t* is the time step of the data generating process. As Δ*t* is typically unknown for empirical data, we have assumed it is equal to 1. multiAR is therefore conceptually similar to multiJI but exploits lag-1 autoregressive models (Hasselmann, 1988) rather than empirical dynamical modelling (Sugihara et al., 2012). Due to this similarity, species selection and data processing was identical to multiJI, with the same expected behaviour; an increase in value with stress.

We should note here, that estimating the dominant real eigenvalue of the Jacobian is only relevant to stable systems not experiencing repeated perturbation(s) (Clark et al., 2024). This is because when no stable equilibrium state is present, or a system is driven by short term rather than long term dynamics, Jacobian estimators are biased. Consequently, these metrics would be appropriate for data from the stable and stressed regions of Figure 1a but not the unstable region. Decision makers interested in stability metrics should therefore believe their community of interest has a) a stable attractor, and b) dynamics are relatively slow or perturbations are infrequent/gradual.

### 2.4 Stability metric trends

All five metrics are expected to express directional change when the system is linearly forced/stressed but remain approximately constant when unstressed (Figure S1). We therefore assessed the linear trend of each metric in a Bayesian mixed effects model as an alternative to the Kendall tau correlation comparisons typically applied to such indicator comparisons (Baruah et al., 2022; Clements et al., 2015; Dutta et al., 2018). Comparing stability metric ability in this way allows us to extract representative global trends of the stochastically simulated communities while accounting for autocorrelation and shared features between simulations (Johnson et al., 2024). Each metric’s trend across time series lengths and search efforts was consequently analysed independently using Laplace approximation in INLA (v.23.04.24) (Rue et al., 2009). Interactions were modelled between time (numeric, scaled between 0 and 1 to allow comparability across truncated time series lengths), whether the simulation represents a stressed or unstressed model (factor, two levels), time series length (numeric, divided by 10 to be of equal magnitude as search effort) and search effort (numeric) as fixed effects. Random intercepts were also included for community identifier (factor, 30 levels) along with an autocorrelation function to account for serial dependence within a simulation, and random slopes for each community as each is expected to display the same trend. Separate models were fit for each community size (5, 15, or 25 species) due to computational constraints. Please note that mvi is modelled on the log scale due to sudden non-linearities known to occur in this metric’s trend (Brock and Carpenter, 2006). The resulting model structure was:

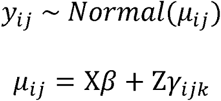

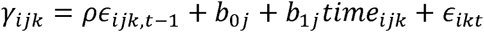

where *y_ij_* is the predicted metric value for the *i*th data point in the *j*th community and *k*th simulation with mean μ*_ij_*. *β* represents the coefficients for time, stress, time series length, search effort and their interactions, *b_0j_* the random intercepts, *b_1j_* the random slope coefficients, *ρ* the correlation between temporally related residuals (*ε*), and *t* the time point of *i*. *X* and *Z* are design matrices for fixed and random effect variables respectively.

We set the weakly informative priors following Rue et al. (2009)’s suggestions:

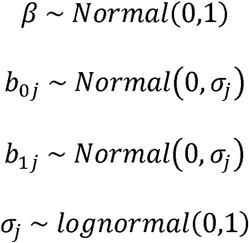

### 2.5 Jacobian threshold transgression

As the Jacobian indices can also signal unstable periods when greater then 1 (for the S-map derived methods) or 0 (for the autoregressive method), we also modelled the capability of these metrics to correctly classify stressed simulations using a binomial mixed effects model. As above, we included time series length, search effort and model stress as fixed effects, with random intercepts for community identifier, but also included community size (numeric) as a fixed effect with the response variable whether the threshold of 1/0 was crossed at any point in the time series, rather than the raw metric value. The model structure was therefore:

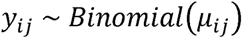

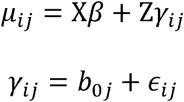

with priors maintained as above.

## 3. Results

### 3.1 Metric trends

All metrics generally display their anticipated behaviour through time and are influenced by both the length of time series and quality of search effort (Figures 2-4, Figures S2-S4). In stressed 5 species communities (Figure 2, Tables S1-S5), increasing search effort improves the strength of trend estimate in all metrics for a given time series length level. However, increasing time series length alters trend differentially between metrics. For example, multiJI’s and FI’s trend is not improved by longer time series while the remainder of metrics all strengthen. Each metric displayed uncertainty in its trend is estimate, owing to the influence of stochasticity in each simulation altering the timing of metric change between simulations and non-linearities in metric trend (Figures S2-S4).

**Figure 2.**
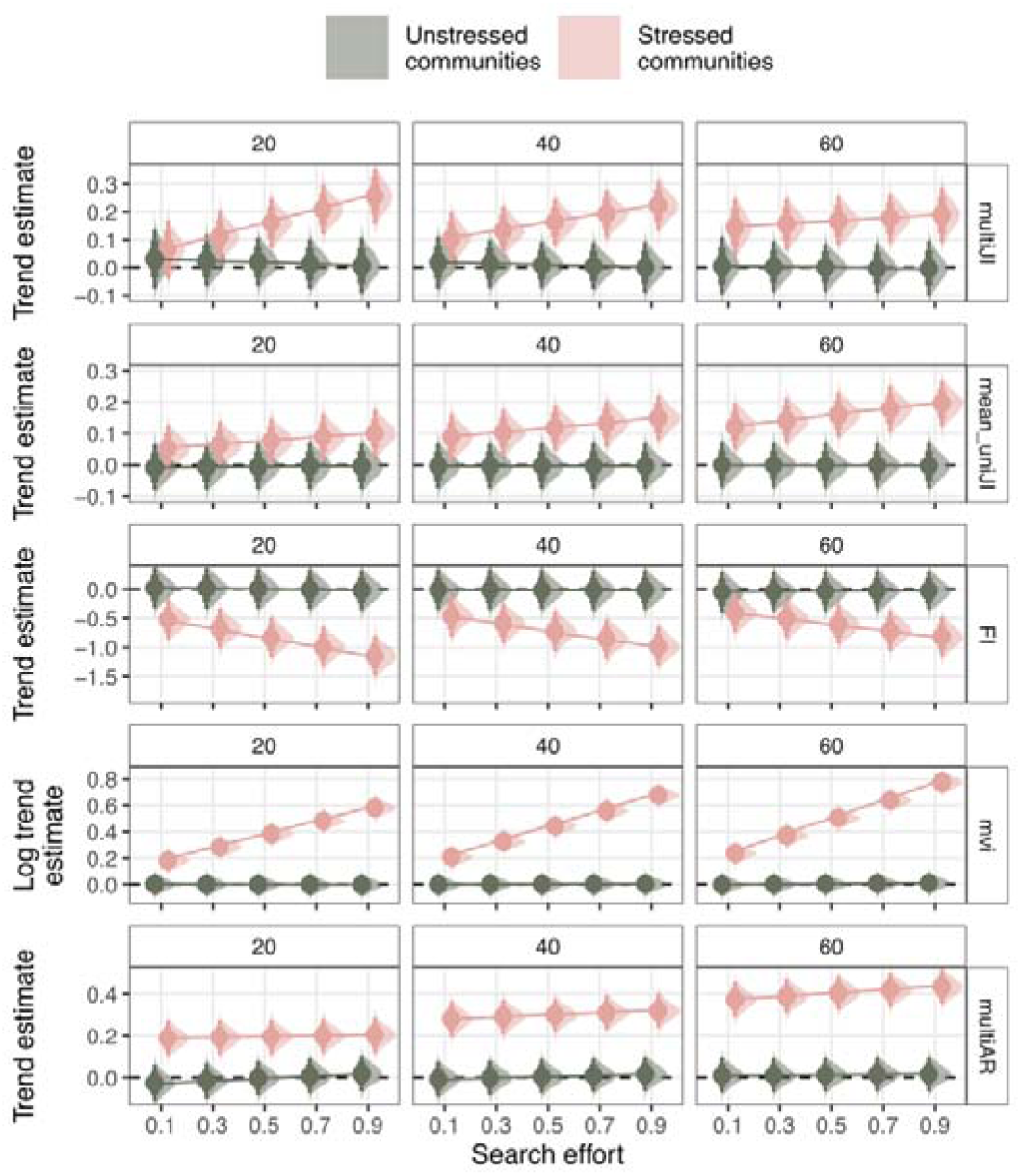
Posterior estimated slopes/trends of each stability metric (rows) through time, across truncated time series lengths (columns) and search efforts in 5 species communities. Dark coloured estimates represent trends in unstressed communities, while light coloured estimates are those in stressed communities. The reported values are the posterior density median values (circles), with 80% (thickest bars) and 95% (thinnest bars) credible intervals.

In the 15 species communities (Figure 3, Tables S6-S10), all metrics other than multiJI and multiAR behaved as anticipated across stressed vs unstressed communities, time series lengths and search efforts: each metric’s trend was approximately zero in the unstressed communities but strengthened with time series length and search effort in the stressed. Both multiJI and mean_uniJI were unreliable in shorter, unstressed communities though became more reliable as time series length increased. However, increasing search effort alters trend differentially between metrics. For example, multiJI’s trend is weakened with improved search efforts while the remainder of metrics all strengthen.

**Figure 3.**
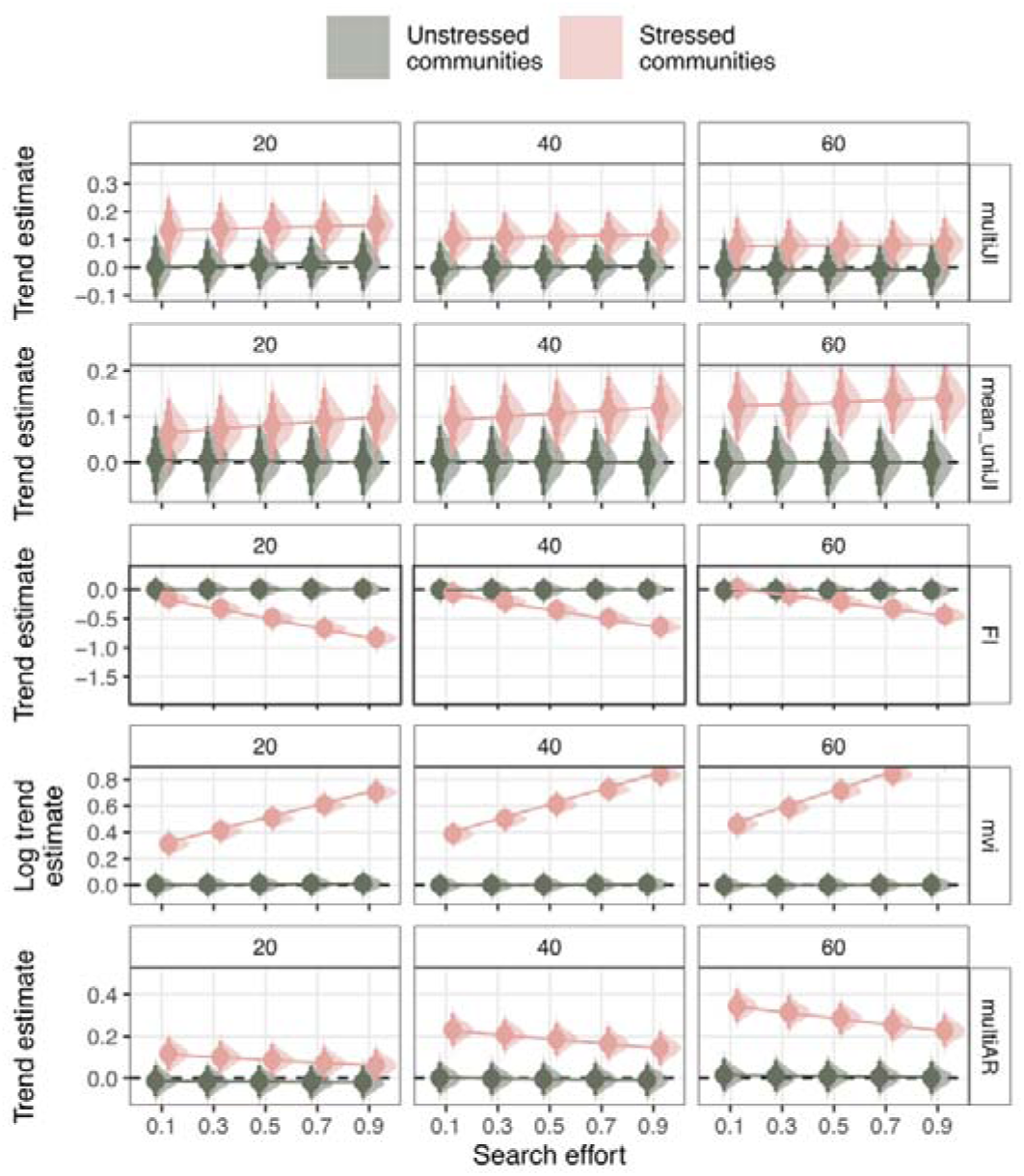
Posterior estimated slopes/trends of each stability metric (rows) through time, across truncated time series lengths (columns) and search efforts in 15 species communities. Dark coloured estimates represent trends in unstressed communities, while light coloured estimates are those in stressed communities. The reported values are the posterior density median values (circles), with 80% (thickest bars) and 95% (thinnest bars) credible intervals.

Finally, all metrics perform less reliably in 25 species communities (Figure 4, Tables S11-S15) relative to the smaller communities. For example, multiJI and mean_uniJI can only distinguish stressed from unstressed in the longest and best searched communities, while both multiJI and multiAR decline in trend as search effort increases. FI and mvi both perform as anticipated across increasing time series lengths and search efforts but with weaker trends than those observed in the smaller communities.

**Figure 4.**
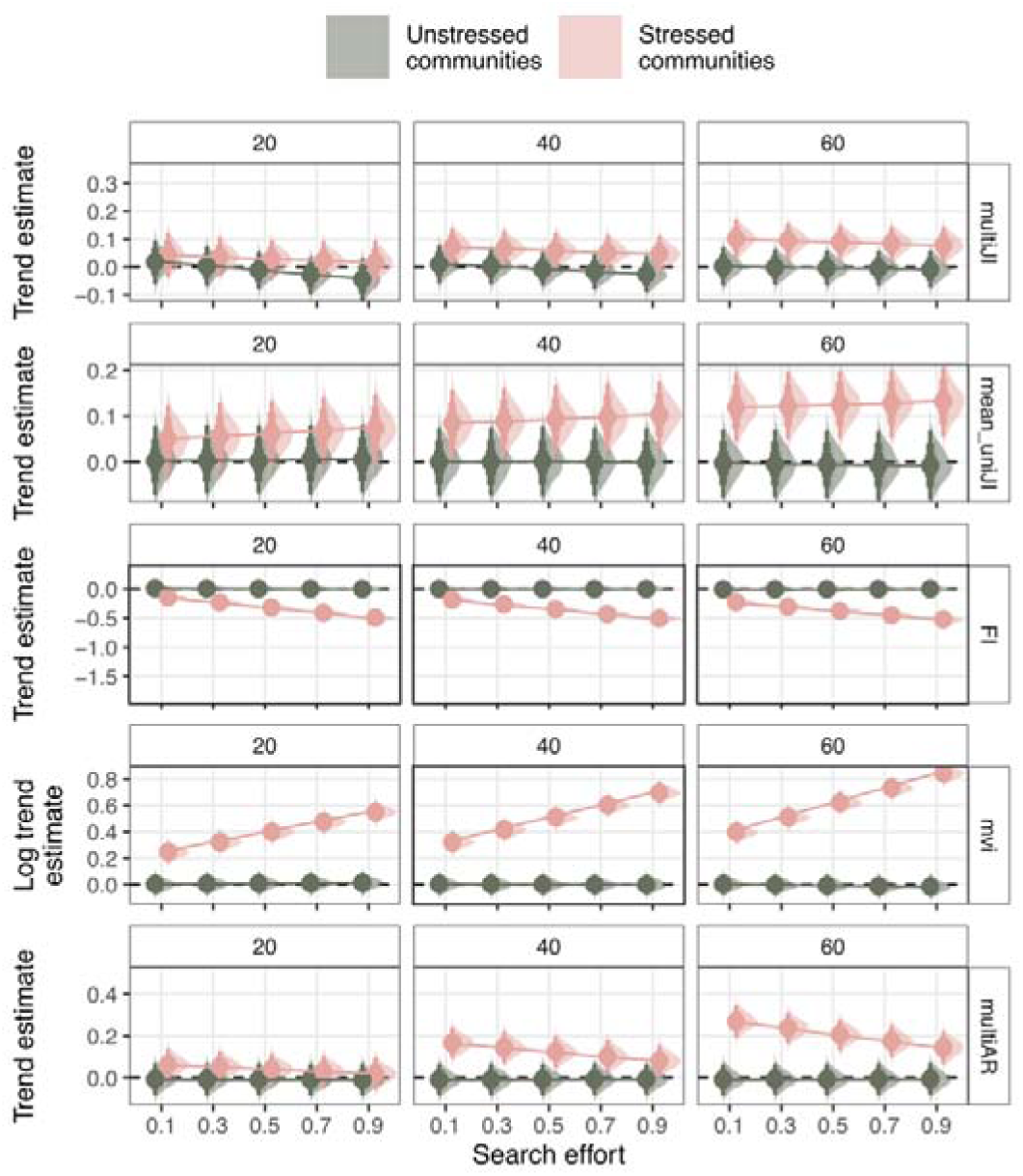
Posterior estimated slopes/trends of each stability metric (rows) through time, across truncated time series lengths (columns) and search efforts in 25 species communities. Dark coloured estimates represent trends in unstressed communities, while light coloured estimates are those in stressed communities. The reported values are the posterior density median values (circles), with 80% (thickest bars) and 95% (thinnest bars) credible intervals.

### 3.2 Threshold transgression

For the Jacobian estimating metrics multiJI, max_uniJI and multiAR, the size of the community and time series length were the best predictors of reported instability probability (Figure 5, Tables S16-S18) – i.e. displaying a metric value greater than its instability threshold. For multiJI and max_uniJI that threshold was 1 and 0 for multiAR. In all metrics, for a constant time series length and search effort, the 25 species community always displayed a higher probability of reported instability than 15 or 5 (Figure 5) while shorter time series also increased the probability of reported instability. max_uniJI always signalled instability regardless of data quality and presence of stress, while multiJI became less likely to report false positives as data quality increased. Search effort did not influence the probability of instability except in multiJI predictions. Time series length generally decreased probability of reported instability but synergised with search effort so that high search effort resulted in equivalent probabilities across time series length.

**Figure 5.**
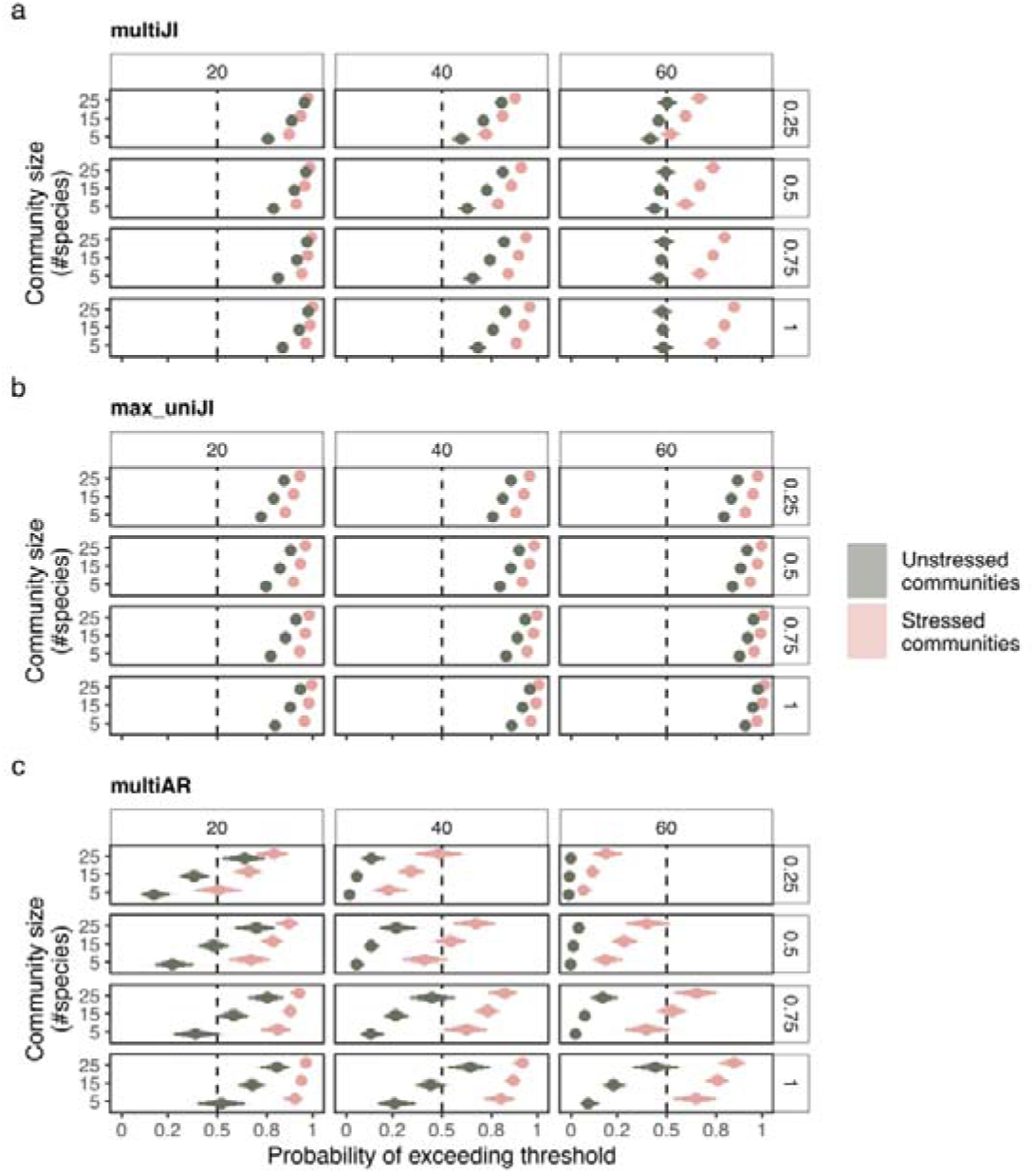
Posterior estimated probabilities of the stability metrics multiJI, max_uniJI and multiAR transgressing its threshold a. Estimates have been delineated by truncated time series lengths (columns) and search efforts (rows) in the three different community sizes. Dark coloured estimates represent trends in unstressed communities, while light coloured estimates are those in stressed communities. The reported values are the posterior density median values (circles), with 80% (thickest bars) and 95% (thinnest bars) credible intervals back transformed from log odds to probabilities.

If the three metrics are compared, multiJI and multiAR displayed identical behaviour in longer time series, implying both estimate similar Jacobians. Overall, max_uniJI displayed both the highest true positive rate (highest probabilities in stressed communities) whereas multiAR displayed the lowest false positive rate (lowest probabilities in unstressed communities). multiJI generally balanced the ratio better than max_uniJI and multiAR with the latter most sensitive to low search efforts. multiJI was also most robust at the highest data qualities (long time series and high search efforts) across community sizes, but more conservative than the other metrics at low data qualities.

## 4. Discussion

Quantifying population and community stability/resilience loss has emerged as a crucial target of ecological research (Kéfi et al., 2019), mapping directly on to the needs of practitioners and policy makers (Donohue et al., 2016; Nicholson et al., 2012). Despite the complexity of ecological systems, political decisions often require simplifications to allow efficient decision making and action (Donatti et al., 2020; Nicholson et al., 2012). Stability metrics represent a bridge between the need for indicators of ecosystem variability, recovery, and simple user interpretation. We show here that stability metrics can perform as anticipated in simulated data analogous to real world biodiversity datasets though the choice of metric and contributing time series requires caution.

### 4.1 Influence of data quality and community size

Our simulations suggest that time series length has a significant impact on their ability to reconstruct estimate stability dimensions as anticipated (Figure S1). In terms of search effort, the multivariate Jacobian metrics (multiJI and multiAR) unexpectedly decreased in reliability as search effort increased. It is unclear why these metrics would behave in this way, though it is possible that, as multiJI and multiAR displayed the highest estimated autocorrelation across metrics (autocorrelation ≈ 1; Tables S1, S5, S6, S10, S11 and S15), the error introduced by low search effort acted as a low pass filter which weakened the influence of autocorrelation and allowed the true interactions between species to be identified. Further, our assumption that search effort linearly influences metric ability may also be violated by the strong non-linearities we observe in the raw indicator trends (Figures S2-S4).

The increasing trend in the metrics with time series length follows the typical behaviour of biodiversity indicators (Clements et al., 2015; Gonzalez et al., 2016; Lenton et al., 2012), with longer time series displaying more reliable estimates of the true trend (Wauchope et al., 2019). This effect is compounded by the caveats of Sugihara et al. (2012) that a minimum of 30 time points are required for reliable empirical dynamic modelling estimates. This is obviously unfeasible for many yearly abundance accounts reported in the LPI and other datasets, and while we show that correct estimates can be made with these short time series, their accuracy is weaker than in time series longer than 30 time points.

It is encouraging, however, that unstressed communities consistently showed no trend in any metric (Figures 2-4) although often signalled instability for metrics with a threshold (Figure 5). This implies that if a trend is observed, then that stability dimension is decreasing prior to abundance decline, and intervention is likely required. Note that no mechanism or driver of stability loss is provided by these metrics and only phenomenological signals are returned. These indicators consequently require system specific knowledge for successful intervention, but this is a recurring challenge for current abundance-based biodiversity indicators such as the LPI (Ledger et al., 2023).

Overall, the dominant determinant of metric performance was the number of species in the simulated community. The multivariate metrics estimating the communities’ Jacobian (multiJI and multiAR) were particularly influenced, with weakening performance as community size increased. Ushio et al. (2018) acknowledge the potential for this degradation and suggest it is dependent on the choice of species included in the Jacobian estimation. We show here that, even when all species contributing to the system are known and measured, including all species compromises the indices’ ability. This is likely due to two considerations. First, increasing the number of species contributing to the stability estimate diffuses the influence of stress through the community due to weak interactions (Clark et al., 2024). This results in the largest communities being more stable compared to small communities. Secondly, the accuracy of the S-map estimated Jacobian is dependent on the state space reconstruction (Ushio et al., 2018). When the number of species in the community increases, the amount of data required to correctly estimate species interactions grows exponentially (Chang et al., 2017). If large communities are analysed with relatively short time series (akin to the time series lengths we simulate here and available to practitioners), then the nearest neighbour step of state-space reconstruction can be misleading (Chang et al., 2021). Fortunately, the S-map technique is an active area of research with advances to mitigate such challenges when working with specious communities (Chang et al., 2021) and ecosystems with strong measurement error (Esguerra and Munch, 2024). These advances unfortunately do not have accessible software compared to the standard S-map used here, but we anticipate that such software will appear and encourage users to consider them.

The univariate Jacobian metric (mean_uniJI) surprisingly outperformed the multivariate (multiJI and multiAR), though it has been suggested that representative variables can provide sufficient dynamics without masking the signal with other uninformative time series (Grziwotz et al., 2023). This does however partially conflict with Donohue et al. (2016, 2013) and Kéfi et al. (2019) who clearly highlight that both uni- and multidimensional measurements of stability are required. To clarify, we agree that biodiversity and indicators need to be multidimensional, but caution against naïve species selection. Appropriate species may be selected based upon their dominance in the ecosystem in terms of abundance (Ushio et al., 2018), causal importance (Chang et al., 2021), or inter-species interaction strengths (Medeiros et al., 2022; Medeiros and Saavedra, 2023; White et al., 2020).

### 4.2 A shift towards multivariate data

A general restriction of the LPI specifically is its primarily single species focus (Ledger et al., 2023; Loh et al., 2005). BIOTIME (Dornelas et al., 2018) and Global Population Dynamics Database (Inchausti and Halley, 2001) have greater assemblage contributions. That said, it is clear that biodiversity monitoring requires multidimensional representation going forward (Donohue et al., 2013; Kéfi et al., 2019; Medeiros et al., 2022; Medeiros and Saavedra, 2023; White et al., 2020) to provide the reliable estimates required for effective governance. Such data will become more readily available with the advent of autonomous and remote ecological monitoring (Besson et al., 2022), but there will be a need for tools capable of synthesising that data in to an interpretable form for decision makers. We suggest these stability metrics can contribute to this synthesis in extension of traditional approaches such as species diversity or richness (Inchausti and Halley, 2001; Turney and Buddle, 2016) due to their development specifically focussing on species interactions. However, if multivariate data is not available, this study supports Grziwotz et al (2023)’s findings that univariate Jacobian estimates can also be reliable indicators of system dynamics in the absence of complete monitoring.

### 4.3 Comparison to critical transition indicators

A similar collection of indicators to the metrics discussed here have received extensive research interest in relation to predicting certain forms of regime shift and population collapse (Baruah et al., 2022; Clements et al., 2015; Dakos, 2018; Dakos et al., 2012). These ‘Early Warning Signals’ (EWSs) can be considered a sub-class of stability indicator (Dakos et al., 2015) as they quantify resilience when a system has two alternative stable states and the capability to transition/regime shift between them. With current debates over the commonality of alternative stable states in ecology (Dakos et al., 2015; Davidson et al., 2023) and inconsistent ability in real world data (Burthe et al., 2016; O’Brien et al., 2023a), the benefit of the explored metrics over EWSs is their ability to quantify stability regardless of the presence of multiple stable states (Lyapunov, 1992; Medeiros et al., 2022; Sugihara et al., 2012). With the complex intrinsic and extrinsic interactions driving ecosystem dynamics possibly masking true multiple stability (Hillebrand et al., 2023) or ecosystems themselves experiencing long transients rather than regime shifts (Hastings et al., 2018), the Jacobian metrics discussed here are conceptually capable of accurate estimation under both heuristics.

### 4.4 Trends vs thresholds

A key challenge for designing indicators appropriate for policy is that governance primarily focusses on thresholds (Donohue et al., 2016; United Nations, 2020) as they allow unambiguous interpretation and enforcement. Conversely, trends require sufficient subjectivity that the vague language associated with trend reporting results in broad objective language (United Nations, 2020). Many governments therefore use indicators with set target thresholds for biodiversity rather than considering a continuous gradient of ‘state’ which is suggested to be more appropriate for modern coupled socio-ecological systems (Hillebrand et al., 2023).

This requirement for a threshold is also explored by the LPI. The debates surrounding the LPI and its interpretation (Dornelas et al., 2019; Leung et al., 2020) ultimately revolve around reconciling the variability of local trend directions (winner and loser populations) with the short time series typically available (Gonzalez et al., 2016). Indeed, controlling for co-dependencies between separate time series such as temporal, spatial and phylogenetic autocorrelation increases the uncertainty of trend estimates (Johnson et al., 2024). This indicator/monitoring uncertainty erodes legislators’ ability to make confident governance, interacts with other forms of uncertainty (e.g. political strategies, lobbying effort, pre-existing laws (Dewulf and Biesbroek, 2018)) and leads to broad objectives. The Jacobian estimating metrics we explored minimise this incongruency between trends and thresholds by facilitating both forms of interpretation. Meanwhile, FI and mvi do not express thresholds but their superior ability relative to Jacobian indices in this study implies trends are more reliable monitoring instruments for ecosystem stability than instability thresholds.

Indeed, instability is not necessarily atypical of natural systems. Lake fish communities can display consistent periods of instability though the year (Ushio et al., 2018), Canadian boreal forests are temporarily unstable following historic fires (Héon et al., 2014), and experimental intertidal communities recover (i.e. display transient instability) following single pulse stress (White et al., 2020). In the metrics tested here, we identified that instability is regularly reported for shorter time series – presumably resulting from transient dynamics within the data window - and that a longer period is necessary for the stable systems to be correctly classified as such. Stability metric thresholds are therefore only informative if instability is prolonged.

## 5. Conclusion

Our study demonstrates stability metrics can accurately quantify stability of simulated multispecies communities manipulated into realistic data qualities. Contributing species/time series is the key consideration for these metrics’ functionality and requires ecological knowledge input from the user. Stability is strongly suggested to be under characterised during global biodiversity monitoring (Capdevila et al., 2022; Kéfi et al., 2019) and the metrics tested here have the potential to contribute generic inference from abundance data alone. We consequently anticipate Jacobian estimation techniques to increase in popularity but pre-empt their use in biodiversity datasets by cautioning towards the longest time series possible and ecologically important species.

## Notes

https://doi.org/10.5281/zenodo.13899465

### Competing Interest Statement

The authors have declared no competing interest.

### Summary of Updates

Clarifying paragraphs regarding empirical dynamical modelling developments and definitions for stability/resilience etc. New conceptual Figure 1 explaining the dimensions of stability. New conceptual Figure S1 explaining the calculations of stability metrics

