## Supplementary information for "Stability metrics behave predictably across data qualities but are sensitive to community size"

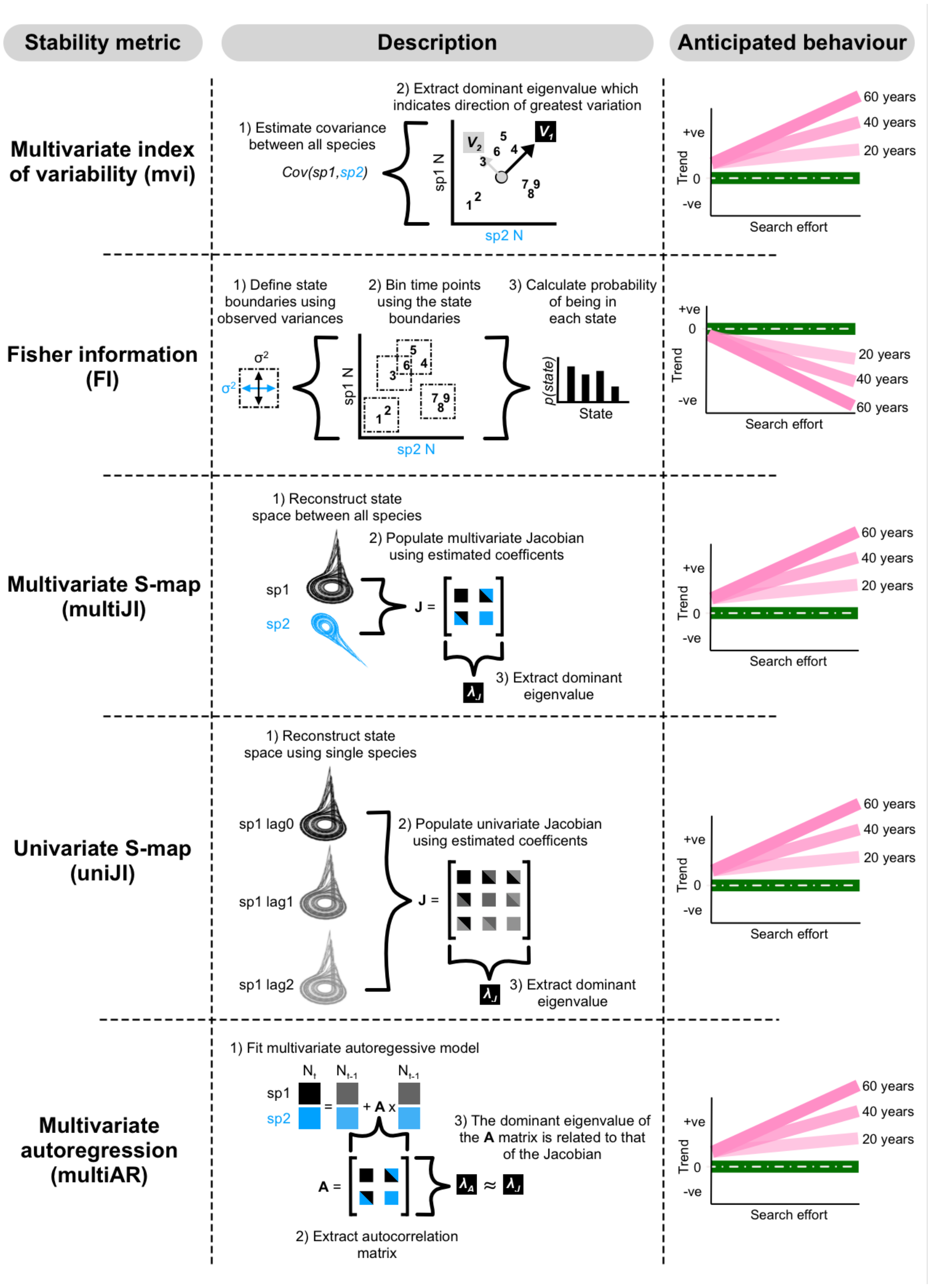

Figure S1 Overview of stability metrics, their calculation and how they are anticipated to behave under unstressed (green) and stressed (pink) scenarios across a range of time series lengths and search efforts.

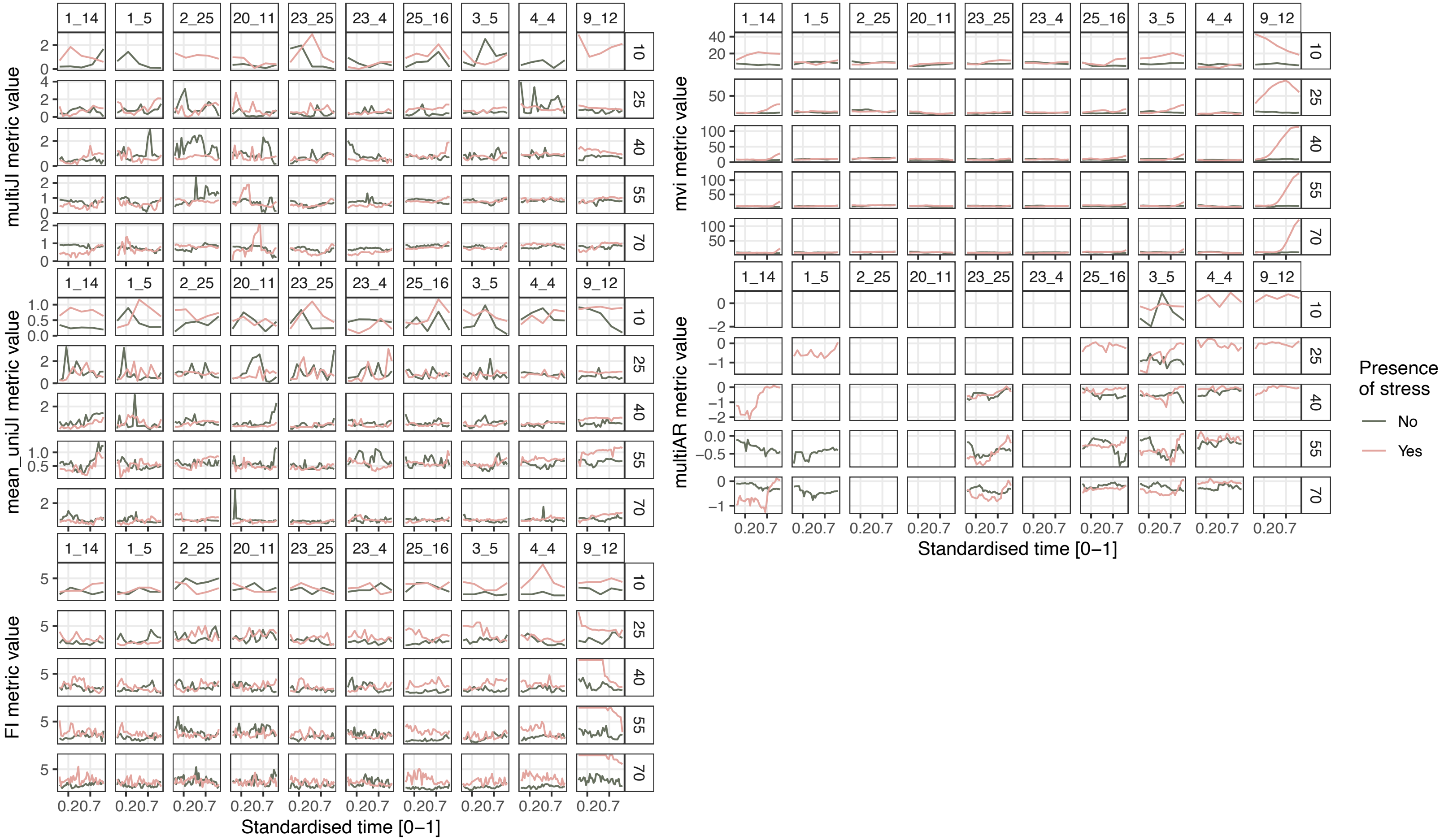

**Figure S2.** Raw stability metric trends in 5 species communities across time series lengths (rows)**.** 10 random simulations (columns) have been selected as examples.

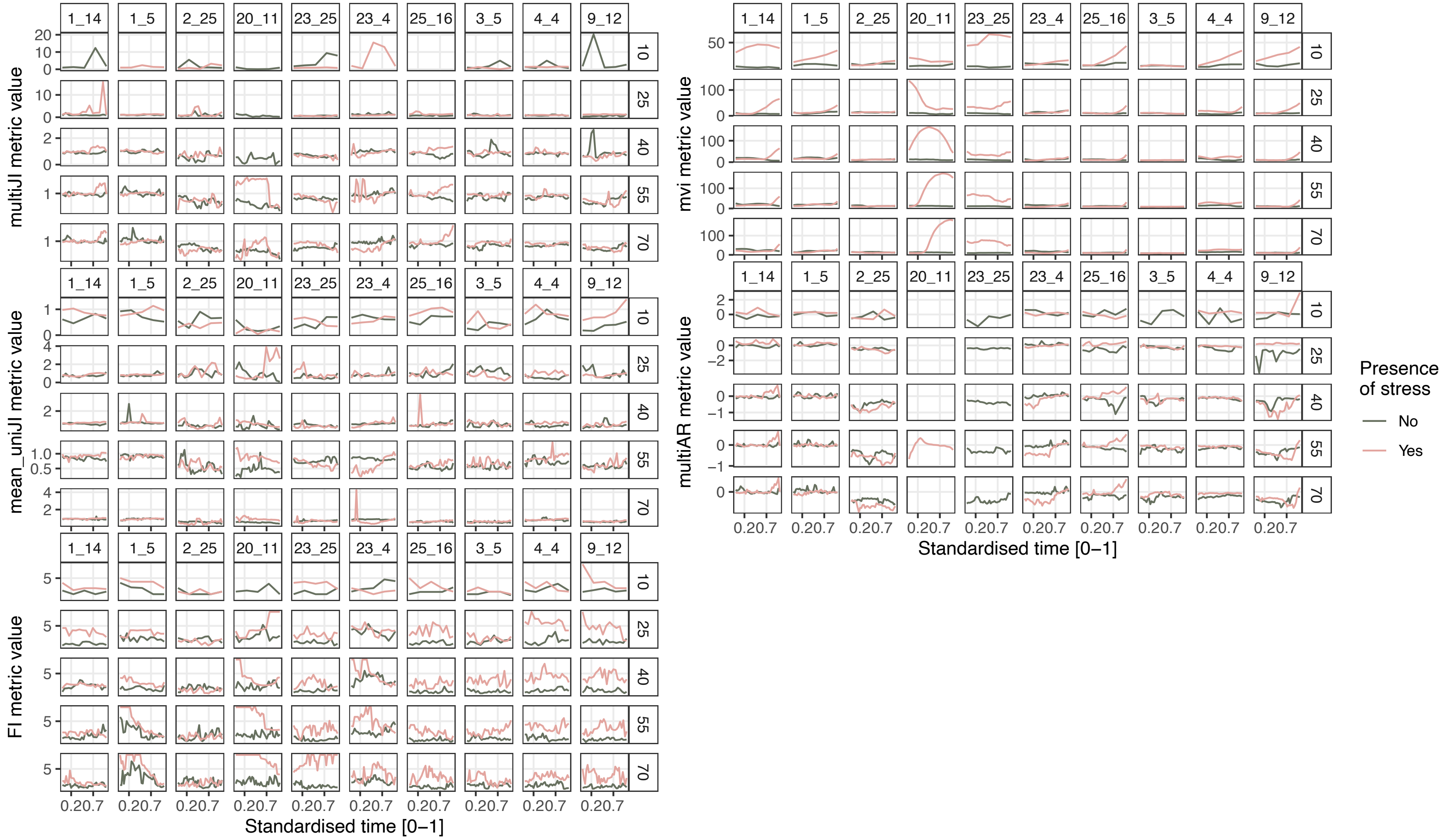

**Figure S3.** Raw stability metric trends in 15 species communities across time series lengths (rows)**.** 10 random simulations (columns) have been selected as examples.

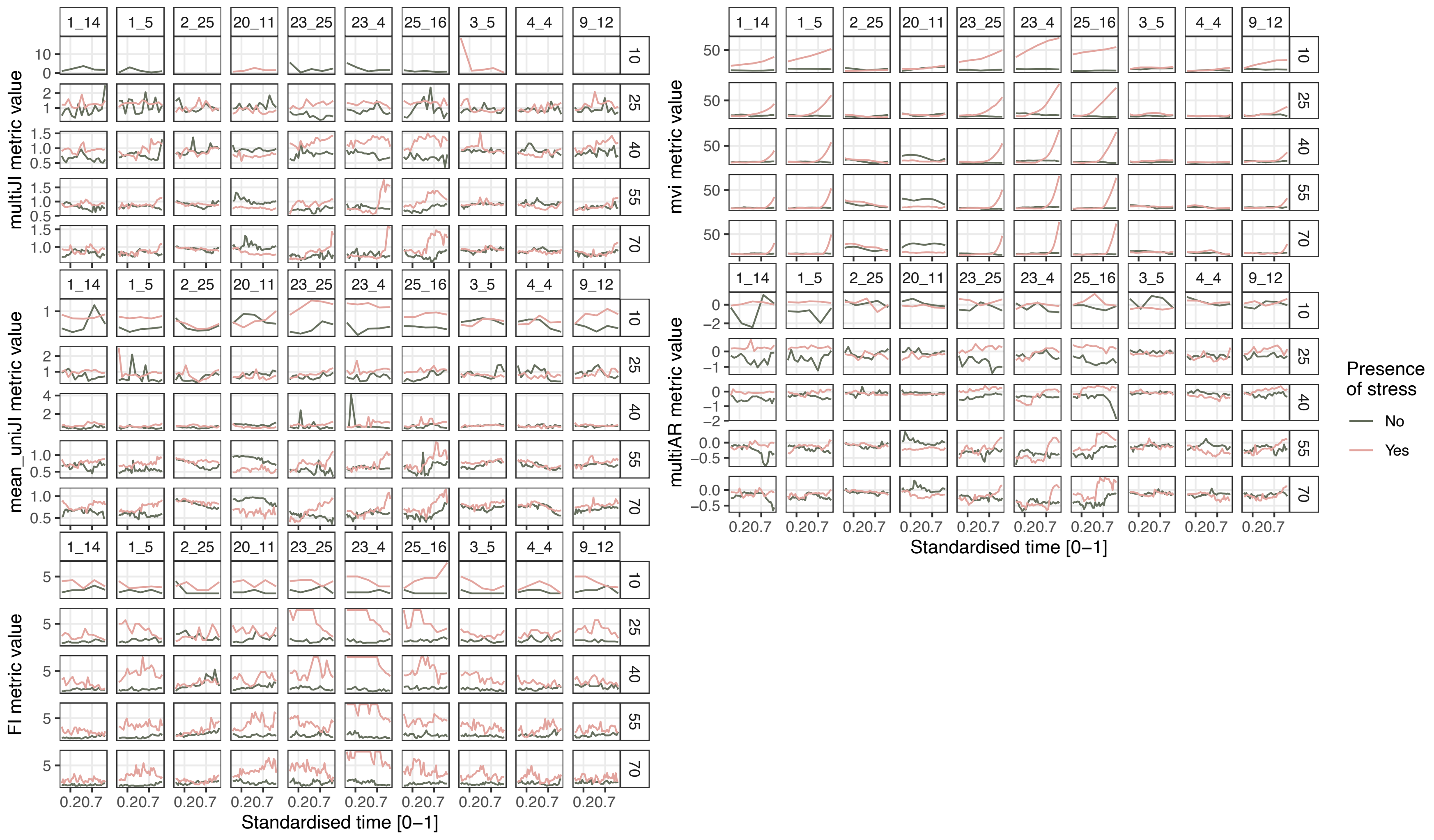

**Figure S4.** Raw stability metric trends in 15 species communities across time series lengths (rows)**.** 10 random simulations columns) have been selected as examples.

**Table S1.** Parameter estimates for multivariate Jacobian index value through time (i.e. stand_time) according to time series length, search effort and stress category in 5 species communities. Median and upper and lower credible intervals are reported.

| Community size | Metric | Parameter | Median | CI_low | CI_high |
|---|---|---|---|---|---|
| 5 | multiJI | *Fixed effects* | | | |
| | | (Intercept) | 1.015 | 0.931 | 1.099 |
| | | stand_time | 0.047 | -0.057 | 0.151 |
| | | stressed1 | 0.344 | 0.254 | 0.433 |
| | | ts_length | -1.27 | -1.405 | -1.135 |
| | | search_effort | 0.083 | -0.019 | 0.185 |
| | | stand_time:stressed1 | -0.057 | -0.203 | 0.088 |
| | | stand_time:ts_length | -0.067 | -0.288 | 0.154 |
| | | stressed1:ts_length | -0.63 | -0.821 | -0.44 |
| | | stand_time:search_effort | -0.038 | -0.205 | 0.129 |
| | | stressed1:search_effort | 0.353 | 0.21 | 0.496 |
| | | ts_length:search_effort | 0.528 | 0.31 | 0.745 |
| | | stand_time:stressed1:ts_length | 0.318 | 0.009 | 0.627 |
| | | stand_time:stressed1:search_effort | 0.372 | 0.14 | 0.605 |
| | | stand_time:ts_length:search_effort | 0.044 | -0.31 | 0.398 |
| | | stressed1:ts_length:search_effort | -0.634 | -0.94 | -0.328 |
| | | stand_time:stressed1:ts_length:search_effort | -0.51 | -1.004 | -0.016 |
| | | *Random effects* | | | |
| | | Precision for comm_id | 50.699 | 50.155 | 51.302 |
| | | Precision for ar_id | 0.298 | 0.296 | 0.301 |
| | | Precision for time | 0.848 | 0.84 | 0.855 |
| | | PACF1 for time | 1 | 1 | 1 |

**Table S2**. Parameter estimates for mean univariate Jacobian index value through time (i.e. stand_time) according to time series length, search effort and stress category in 5 species communities. Median and upper and lower credible intervals are reported.

| Community size | Metric | Parameter | Median | CI_low | CI_high |
|---|---|---|---|---|---|
| 5 | mean_uniJI | *Fixed effects* | | | |
| | | (Intercept) | 0.661 | 0.607 | 0.715 |
| | | stand_time | -0.012 | -0.034 | 0.01 |
| | | stressed1 | 0.012 | -0.01 | 0.035 |
| | | ts_length | -0.55 | -0.582 | -0.518 |
| | | search_effort | 0.097 | 0.071 | 0.122 |
| | | stand_time:stressed1 | 0.027 | -0.004 | 0.059 |
| | | stand_time:ts_length | 0.02 | -0.028 | 0.068 |
| | | stressed1:ts_length | -0.038 | -0.083 | 0.008 |
| | | stand_time:search_effort | 0.009 | -0.027 | 0.045 |
| | | stressed1:search_effort | 0.131 | 0.094 | 0.167 |
| | | ts_length:search_effort | 0.33 | 0.278 | 0.382 |
| | | stand_time:stressed1:ts_length | 0.144 | 0.077 | 0.212 |
| | | stand_time:stressed1:search_effort | 0.033 | -0.018 | 0.084 |
| | | stand_time:ts_length:search_effort | -0.018 | -0.095 | 0.059 |
| | | stressed1:ts_length:search_effort | -0.252 | -0.325 | -0.179 |
| | | stand_time:stressed1:ts_length:search_effort | 0.102 | -0.007 | 0.211 |
| | | *Random effects* | | | |
| | | Precision for comm_id | 57.955 | 57.363 | 58.564 |
| | | Precision for ar_id | 218.473 | 216.257 | 220.817 |
| | | Precision for time | 10.44 | 10.361 | 10.517 |
| | | PACF1 for time | 0.793 | 0.792 | 0.795 |

**Table S3**. Parameter estimates for Fisher information value through time (i.e. stand_time) according to time series length, search effort and stress category in 5 species communities. Median and upper and lower credible intervals are reported.

| Community size | Metric | Parameter | Median | CI_low | CI_high |
|---|---|---|---|---|---|
| 5 | FI | *Fixed effects* | | | |
| | | (Intercept) | 2.308 | 2.059 | 2.556 |
| | | stand_time | 0.064 | 0.009 | 0.118 |
| | | stressed1 | 0.582 | 0.503 | 0.66 |
| | | ts_length | -0.216 | -0.336 | -0.096 |
| | | search_effort | -0.026 | -0.116 | 0.064 |
| | | stand_time:stressed1 | -0.553 | -0.629 | -0.476 |
| | | stand_time:ts_length | -0.192 | -0.324 | -0.06 |
| | | stressed1:ts_length | 0.811 | 0.643 | 0.98 |
| | | stand_time:search_effort | -0.087 | -0.175 | 0 |
| | | stressed1:search_effort | 1.165 | 1.039 | 1.291 |
| | | ts_length:search_effort | -0.393 | -0.586 | -0.199 |
| | | stand_time:stressed1:ts_length | 0.431 | 0.246 | 0.617 |
| | | stand_time:stressed1:search_effort | -0.823 | -0.946 | -0.699 |
| | | stand_time:ts_length:search_effort | 0.2 | -0.013 | 0.413 |
| | | stressed1:ts_length:search_effort | -0.229 | -0.5 | 0.042 |
| | | stand_time:stressed1:ts_length:search_effort | 0.444 | 0.146 | 0.742 |
| | | *Random effects* | | | |
| | | Precision for comm_id | 2.572 | 2.526 | 2.616 |
| | | Precision for ar_id | 41.051 | 39.39 | 42.547 |
| | | Precision for time | 0.991 | 0.98 | 1.003 |
| | | PACF1 for time | 0.96 | 0.96 | 0.96 |

**Table S4**. Parameter estimates for multivariate index of variability value through time (i.e. stand_time) according to time series length, search effort and stress category in 5 species communities. Median and upper and lower credible intervals are reported.

| Community size | Metric | Parameter | Median | CI_low | CI_high |
|---|---|---|---|---|---|
| 5 | mvi | *Fixed effects* | | | |
| | | (Intercept) | 1.356 | 1.3 | 1.412 |
| | | stand_time | 0.005 | -0.015 | 0.025 |
| | | stressed1 | 0.06 | 0.036 | 0.084 |
| | | ts_length | -0.013 | -0.05 | 0.024 |
| | | search_effort | 0.846 | 0.819 | 0.873 |
| | | stand_time:stressed1 | 0.11 | 0.081 | 0.138 |
| | | stand_time:ts_length | -0.013 | -0.061 | 0.036 |
| | | stressed1:ts_length | -0.054 | -0.106 | -0.001 |
| | | stand_time:search_effort | -0.016 | -0.048 | 0.017 |
| | | stressed1:search_effort | 0.324 | 0.285 | 0.362 |
| | | ts_length:search_effort | 0.319 | 0.259 | 0.379 |
| | | stand_time:stressed1:ts_length | 0.104 | 0.035 | 0.172 |
| | | stand_time:stressed1:search_effort | 0.43 | 0.384 | 0.476 |
| | | stand_time:ts_length:search_effort | 0.051 | -0.027 | 0.129 |
| | | stressed1:ts_length:search_effort | -0.609 | -0.693 | -0.524 |
| | | stand_time:stressed1:ts_length:search_effort | 0.377 | 0.267 | 0.488 |
| | | *Random effects* | | | |
| | | Precision for comm_id | 54.514 | 53.022 | 56.206 |
| | | Precision for ar_id | 8.655 | 8.575 | 8.761 |
| | | Precision for time | 9.065 | 9.038 | 9.107 |
| | | PACF1 for time | 0.976 | 0.975 | 0.976 |

**Table S5.** Parameter estimates for multivariate autocorrelation value through time (i.e. stand_time) according to time series length, search effort and stress category in 5 species communities. Median and upper and lower credible intervals are reported.

| Community size | Metric | Parameter | Median | CI_low | CI_high |
|---|---|---|---|---|---|
| 5 | multiAR | *Fixed effects* | | | |
| | | (Intercept) | -0.632 | -0.702 | -0.563 |
| | | stand_time | -0.054 | -0.104 | -0.003 |
| | | stressed1 | 0.103 | 0.052 | 0.154 |
| | | ts_length | -1.529 | -1.606 | -1.452 |
| | | search_effort | -0.018 | -0.082 | 0.046 |
| | | stand_time:stressed1 | 0.152 | 0.088 | 0.216 |
| | | stand_time:ts_length | 0.107 | 0.001 | 0.213 |
| | | stressed1:ts_length | -0.283 | -0.386 | -0.181 |
| | | stand_time:search_effort | 0.082 | 0 | 0.164 |
| | | stressed1:search_effort | 0.4 | 0.318 | 0.482 |
| | | ts_length:search_effort | 1.848 | 1.723 | 1.974 |
| | | stand_time:stressed1:ts_length | 0.34 | 0.199 | 0.48 |
| | | stand_time:stressed1:search_effort | -0.092 | -0.196 | 0.011 |
| | | stand_time:ts_length:search_effort | -0.123 | -0.296 | 0.049 |
| | | stressed1:ts_length:search_effort | -0.685 | -0.851 | -0.52 |
| | | stand_time:stressed1:ts_length:search_effort | 0.27 | 0.044 | 0.496 |
| | | *Random effects* | | | |
| | | Precision for comm_id | 53.967 | 50.41 | 57.52 |
| | | Precision for ar_id | 29.391 | 26.573 | 32.763 |
| | | Precision for time | 4.778 | 4.745 | 4.811 |
| | | PACF1 for time | 0.884 | 0.883 | 0.885 |

**Table S6.** Parameter estimates for multivariate Jacobian index value through time (i.e. stand_time) according to time series length, search effort and stress category in 15 species communities. Median and upper and lower credible intervals are reported.

| Community size | Metric | Parameter | Median | CI_low | CI_high |
|---|---|---|---|---|---|
| 15 | multiJI | *Fixed effects* | | | |
| | | (Intercept) | 1.436 | 1.341 | 1.531 |
| | | stand_time | -0.001 | -0.11 | 0.108 |
| | | stressed1 | -0.032 | -0.146 | 0.082 |
| | | ts_length | -1.657 | -1.822 | -1.493 |
| | | search_effort | 0.383 | 0.256 | 0.51 |
| | | stand_time:stressed1 | 0.159 | 0.003 | 0.315 |
| | | stand_time:ts_length | -0.004 | -0.228 | 0.22 |
| | | stressed1:ts_length | 0.102 | -0.133 | 0.337 |
| | | stand_time:search_effort | 0.043 | -0.132 | 0.217 |
| | | stressed1:search_effort | 0.14 | -0.043 | 0.324 |
| | | ts_length:search_effort | -0.112 | -0.376 | 0.152 |
| | | stand_time:stressed1:ts_length | -0.13 | -0.449 | 0.189 |
| | | stand_time:stressed1:search_effort | -0.011 | -0.261 | 0.24 |
| | | stand_time:ts_length:search_effort | -0.082 | -0.441 | 0.277 |
| | | stressed1:ts_length:search_effort | -0.207 | -0.584 | 0.171 |
| | | stand_time:stressed1:ts_length:search_effort | 0.042 | -0.469 | 0.552 |
| | | *Random effects* | | | |
| | | Precision for comm_id | 54.758 | 51.331 | 57.855 |
| | | Precision for ar_id | 0.345 | 0.34 | 0.353 |
| | | Precision for time | 0.628 | 0.618 | 0.639 |
| | | PACF1 for time | 1 | 1 | 1 |

**Table S7.** Parameter estimates for mean univariate Jacobian index value through time (i.e. stand_time) according to time series length, search effort and stress category in 15 species communities. Median and upper and lower credible intervals are reported.

| Community size | Metric | Parameter | Median | CI_low | CI_high |
|---|---|---|---|---|---|
| 15 | mean_uniJI | *Fixed effects* | | | |
| | | (Intercept) | 0.594 | 0.54 | 0.648 |
| | | stand_time | 0.006 | -0.01 | 0.023 |
| | | stressed1 | 0.022 | 0.006 | 0.039 |
| | | ts_length | -0.516 | -0.539 | -0.492 |
| | | search_effort | 0.071 | 0.052 | 0.089 |
| | | stand_time:stressed1 | 0.022 | -0.002 | 0.045 |
| | | stand_time:ts_length | -0.008 | -0.044 | 0.029 |
| | | stressed1:ts_length | -0.003 | -0.037 | 0.031 |
| | | stand_time:search_effort | -0.005 | -0.032 | 0.022 |
| | | stressed1:search_effort | 0.137 | 0.111 | 0.164 |
| | | ts_length:search_effort | 0.561 | 0.523 | 0.599 |
| | | stand_time:stressed1:ts_length | 0.161 | 0.109 | 0.213 |
| | | stand_time:stressed1:search_effort | 0.058 | 0.02 | 0.097 |
| | | stand_time:ts_length:search_effort | 0.003 | -0.056 | 0.062 |
| | | stressed1:ts_length:search_effort | -0.256 | -0.31 | -0.202 |
| | | stand_time:stressed1:ts_length:search_effort | -0.055 | -0.138 | 0.028 |
| | | *Random effects* | | | |
| | | Precision for comm_id | 55.513 | 54.932 | 56.24 |
| | | Precision for ar_id | 85.259 | 83.662 | 87.363 |
| | | Precision for time | 22.032 | 21.812 | 22.199 |
| | | PACF1 for time | 0.825 | 0.824 | 0.827 |

**Table S8.** Parameter estimates for Fisher information value through time (i.e. stand_time) according to time series length, search effort and stress category in 15 species communities. Median and upper and lower credible intervals are reported.

| Community size | Metric | Parameter | Median | CI_low | CI_high |
|---|---|---|---|---|---|
| 15 | FI | *Fixed effects* | | | |
| | | (Intercept) | 1.575 | 1.437 | 1.712 |
| | | stand_time | 0.004 | -0.056 | 0.063 |
| | | stressed1 | 1.316 | 1.239 | 1.393 |
| | | ts_length | -0.516 | -0.633 | -0.399 |
| | | search_effort | 0.27 | 0.183 | 0.357 |
| | | stand_time:stressed1 | -0.148 | -0.232 | -0.064 |
| | | stand_time:ts_length | -0.026 | -0.169 | 0.117 |
| | | stressed1:ts_length | 1.087 | 0.922 | 1.252 |
| | | stand_time:search_effort | 0.013 | -0.083 | 0.108 |
| | | stressed1:search_effort | 1.117 | 0.994 | 1.241 |
| | | ts_length:search_effort | -0.093 | -0.281 | 0.095 |
| | | stand_time:stressed1:ts_length | 0.399 | 0.197 | 0.6 |
| | | stand_time:stressed1:search_effort | -0.994 | -1.129 | -0.859 |
| | | stand_time:ts_length:search_effort | -0.026 | -0.255 | 0.204 |
| | | stressed1:ts_length:search_effort | -0.992 | -1.258 | -0.727 |
| | | stand_time:stressed1:ts_length:search_effort | 0.701 | 0.378 | 1.025 |
| | | Precision for comm_id | 9.42 | 7.758 | 11.121 |
| | | Precision for ar_id | 3.352 | 3.265 | 3.459 |
| | | Precision for time | 0.975 | 0.965 | 0.984 |
| | | PACF1 for time | 0.953 | 0.952 | 0.953 |

**Table S9.** Parameter estimates for multivariate index of variability value through time (i.e. stand_time) according to time series length, search effort and stress category in 15 species communities. Median and upper and lower credible intervals are reported.

| **Community size** | **Metric** | **Parameter** | **Median** | **CI_low** | **CI_high** |
|---|---|---|---|---|---|
| 15 | mvi | *Fixed effects* | | | |
| | | (Intercept) | 1.382 | 1.32 | 1.445 |
| | | stand_time | 0.001 | -0.022 | 0.024 |
| | | stressed1 | 0.145 | 0.114 | 0.176 |
| | | ts_length | -0.187 | -0.236 | -0.139 |
| | | search_effort | 0.834 | 0.798 | 0.869 |
| | | stand_time:stressed1 | 0.199 | 0.167 | 0.231 |
| | | stand_time:ts_length | -0.01 | -0.064 | 0.044 |
| | | stressed1:ts_length | -0.044 | -0.112 | 0.025 |
| | | stand_time:search_effort | 0.015 | -0.021 | 0.052 |
| | | stressed1:search_effort | 0.511 | 0.461 | 0.561 |
| | | ts_length:search_effort | 0.726 | 0.648 | 0.804 |
| | | stand_time:stressed1:ts_length | 0.335 | 0.258 | 0.412 |
| | | stand_time:stressed1:search_effort | 0.4 | 0.348 | 0.451 |
| | | stand_time:ts_length:search_effort | -0.012 | -0.1 | 0.075 |
| | | stressed1:ts_length:search_effort | -0.761 | -0.872 | -0.651 |
| | | stand_time:stressed1:ts_length:search_effort | 0.384 | 0.26 | 0.507 |
| | | *Random effects* | | | |
| | | Precision for comm_id | 45.236 | 44.89 | 45.63 |
| | | Precision for ar_id | 6.754 | 6.708 | 6.799 |
| | | Precision for time | 5.341 | 5.324 | 5.358 |
| | | PACF1 for time | 0.985 | 0.984 | 0.985 |

**Table S10**. Parameter estimates for multivariate autocorrelation value through time (i.e. stand_time) according to time series length, search effort and stress category in 15 species communities. Median and upper and lower credible intervals are reported.

| **Community size** | **Metric** | **Parameter** | **Median** | **CI_low** | **CI_high** |
|---|---|---|---|---|---|
| 15 | multiAR | *Fixed effects* | | | |
| | | (Intercept) | -0.282 | -0.338 | -0.226 |
| | | stand_time | -0.029 | -0.052 | -0.005 |
| | | stressed1 | 0.136 | 0.109 | 0.162 |
| | | ts_length | -1.401 | -1.439 | -1.362 |
| | | search_effort | 0.046 | 0.017 | 0.076 |
| | | stand_time:stressed1 | 0.032 | -0.002 | 0.067 |
| | | stand_time:ts_length | 0.079 | 0.023 | 0.134 |
| | | stressed1:ts_length | -0.216 | -0.272 | -0.16 |
| | | stand_time:search_effort | 0 | -0.038 | 0.037 |
| | | stressed1:search_effort | 0.242 | 0.199 | 0.284 |
| | | ts_length:search_effort | 1.439 | 1.377 | 1.502 |
| | | stand_time:stressed1:ts_length | 0.517 | 0.437 | 0.597 |
| | | stand_time:stressed1:search_effort | -0.025 | -0.08 | 0.029 |
| | | stand_time:ts_length:search_effort | -0.028 | -0.117 | 0.061 |
| | | stressed1:ts_length:search_effort | -0.375 | -0.465 | -0.285 |
| | | stand_time:stressed1:ts_length:search_effort | -0.174 | -0.302 | -0.046 |
| | | *Random effects* | | | |
| | | Precision for comm_id | 54.326 | 53.617 | 55.079 |
| | | Precision for ar_id | 15.618 | 15.468 | 15.767 |
| | | Precision for time | 9.152 | 9.068 | 9.239 |
| | | PACF1 for time | 0.92 | 0.92 | 0.921 |

**Table S11.** Parameter estimates for multivariate Jacobian index value through time (i.e. stand_time) according to time series length, search effort and stress category in 25 species communities. Median and upper and lower credible intervals are reported.

| Community size | Metric | Parameter | Median | CI_low | CI_high |
|---|---|---|---|---|---|
| 25 | multiJI | *Fixed effects* | | | |
| | | (Intercept) | 1.296 | 1.217 | 1.375 |
| | | stand_time | 0.038 | -0.046 | 0.123 |
| | | stressed1 | 0.239 | 0.144 | 0.334 |
| | | ts_length | -1.32 | -1.45 | -1.19 |
| | | search_effort | 0.367 | 0.26 | 0.474 |
| | | stand_time:stressed1 | -0.02 | -0.14 | 0.099 |
| | | stand_time:ts_length | -0.056 | -0.212 | 0.099 |
| | | stressed1:ts_length | -0.324 | -0.508 | -0.14 |
| | | stand_time:search_effort | -0.112 | -0.247 | 0.023 |
| | | stressed1:search_effort | 0.021 | -0.131 | 0.174 |
| | | ts_length:search_effort | -0.088 | -0.296 | 0.121 |
| | | stand_time:stressed1:ts_length | 0.201 | -0.019 | 0.421 |
| | | stand_time:stressed1:search_effort | 0.075 | -0.117 | 0.267 |
| | | stand_time:ts_length:search_effort | 0.164 | -0.085 | 0.413 |
| | | stressed1:ts_length:search_effort | -0.029 | -0.325 | 0.267 |
| | | stand_time:stressed1:ts_length:search_effort | -0.15 | -0.503 | 0.204 |
| | | *Random effects* | | | |
| | | Precision for comm_id | 88.251 | 86.091 | 90.483 |
| | | Precision for ar_id | 1930.483 | 1897.68 | 1963.984 |
| | | Precision for time | 2.072 | 2.028 | 2.117 |
| | | PACF1 for time | 1 | 1 | 1 |

**Table S12.** Parameter estimates for mean univariate Jacobian index value through time (i.e. stand_time) according to time series length, search effort and stress category in 25 species communities. Median and upper and lower credible intervals are reported.

| Community size | Metric | Parameter | Median | CI_low | CI_high |
|---|---|---|---|---|---|
| 25 | mean_uniJI | *Fixed effects* | | | |
| | | (Intercept) | 0.584 | 0.53 | 0.637 |
| | | stand_time | 0.003 | -0.012 | 0.018 |
| | | stressed1 | 0 | -0.016 | 0.015 |
| | | ts_length | -0.512 | -0.535 | -0.489 |
| | | search_effort | 0.051 | 0.034 | 0.069 |
| | | stand_time:stressed1 | 0.008 | -0.013 | 0.029 |
| | | stand_time:ts_length | -0.009 | -0.042 | 0.024 |
| | | stressed1:ts_length | 0.034 | 0.002 | 0.066 |
| | | stand_time:search_effort | 0.011 | -0.013 | 0.035 |
| | | stressed1:search_effort | 0.148 | 0.122 | 0.173 |
| | | ts_length:search_effort | 0.605 | 0.568 | 0.641 |
| | | stand_time:stressed1:ts_length | 0.187 | 0.14 | 0.234 |
| | | stand_time:stressed1:search_effort | 0.026 | -0.008 | 0.06 |
| | | stand_time:ts_length:search_effort | -0.03 | -0.082 | 0.023 |
| | | stressed1:ts_length:search_effort | -0.249 | -0.301 | -0.197 |
| | | stand_time:stressed1:ts_length:search_effort | -0.008 | -0.083 | 0.067 |
| | | *Random effects* | | | |
| | | Precision for comm_id | 55.198 | 54.411 | 55.999 |
| | | Precision for ar_id | 290.896 | 285.242 | 296.717 |
| | | Precision for time | 25.777 | 25.515 | 26.038 |
| | | PACF1 for time | 0.859 | 0.857 | 0.86 |

**Table S13.** Parameter estimates for Fisher information value through time (i.e. stand_time) according to time series length, search effort and stress category in 25 species communities. Median and upper and lower credible intervals are reported.

| Community size | Metric | Parameter | Median | CI_low | CI_high |
|---|---|---|---|---|---|
| 25 | FI | *Fixed effects* | | | |
| | | (Intercept) | 1.542 | 1.465 | 1.62 |
| | | stand_time | 0.016 | -0.04 | 0.072 |
| | | stressed1 | 1.192 | 1.118 | 1.266 |
| | | ts_length | -0.806 | -0.918 | -0.694 |
| | | search_effort | 0.099 | 0.016 | 0.183 |
| | | stand_time:stressed1 | -0.068 | -0.147 | 0.012 |
| | | stand_time:ts_length | -0.044 | -0.179 | 0.091 |
| | | stressed1:ts_length | 1.941 | 1.783 | 2.1 |
| | | stand_time:search_effort | -0.023 | -0.113 | 0.067 |
| | | stressed1:search_effort | 0.842 | 0.723 | 0.96 |
| | | ts_length:search_effort | 0.158 | -0.022 | 0.339 |
| | | stand_time:stressed1:ts_length | -0.193 | -0.384 | -0.002 |
| | | stand_time:stressed1:search_effort | -0.455 | -0.582 | -0.327 |
| | | stand_time:ts_length:search_effort | 0.055 | -0.163 | 0.272 |
| | | stressed1:ts_length:search_effort | -1.075 | -1.33 | -0.82 |
| | | stand_time:stressed1:ts_length:search_effort | 0.132 | -0.175 | 0.439 |
| | | *Random effects* | | | |
| | | Precision for comm_id | 47.01 | 46.609 | 47.417 |
| | | Precision for ar_id | 3.961 | 3.851 | 4.079 |
| | | Precision for time | 1.068 | 1.059 | 1.079 |
| | | PACF1 for time | 0.954 | 0.954 | 0.955 |

**Table S14.** Parameter estimates for multivariate index of variability value through time (i.e. stand_time) according to time series length, search effort and stress category in 15 species communities. Median and upper and lower credible intervals are reported.

| Community size | Metric | Parameter | Median | CI_low | CI_high |
|---|---|---|---|---|---|
| 25 | mvi | *Fixed effects* | | | |
| | | (Intercept) | 1.397 | 1.34 | 1.454 |
| | | stand_time | -0.002 | -0.025 | 0.021 |
| | | stressed1 | 0.107 | 0.077 | 0.137 |
| | | ts_length | -0.205 | -0.252 | -0.159 |
| | | search_effort | 0.837 | 0.803 | 0.871 |
| | | stand_time:stressed1 | 0.142 | 0.109 | 0.175 |
| | | stand_time:ts_length | 0.011 | -0.045 | 0.066 |
| | | stressed1:ts_length | 0.014 | -0.052 | 0.08 |
| | | stand_time:search_effort | 0.033 | -0.004 | 0.071 |
| | | stressed1:search_effort | 0.487 | 0.439 | 0.535 |
| | | ts_length:search_effort | 0.762 | 0.687 | 0.836 |
| | | stand_time:stressed1:ts_length | 0.327 | 0.249 | 0.406 |
| | | stand_time:stressed1:search_effort | 0.262 | 0.208 | 0.315 |
| | | stand_time:ts_length:search_effort | -0.094 | -0.183 | -0.005 |
| | | stressed1:ts_length:search_effort | -0.654 | -0.76 | -0.548 |
| | | stand_time:stressed1:ts_length:search_effort | 0.528 | 0.401 | 0.654 |
| | | *Random effects* | | | |
| | | Precision for comm_id | 55.441 | 54.871 | 56.067 |
| | | Precision for ar_id | 6.061 | 6.012 | 6.11 |
| | | Precision for time | 5.836 | 5.808 | 5.865 |
| | | PACF1 for time | 0.985 | 0.985 | 0.985 |

**Table S15.** Parameter estimates for multivariate autocorrelation value through time (i.e. stand_time) according to time series length, search effort and stress category in 25 species communities. Median and upper and lower credible intervals are reported.

| Community size | Metric | Parameter | Median | CI_low | CI_high |
|---|---|---|---|---|---|
| 25 | multiAR | *Fixed effects* | | | |
| | | (Intercept) | -0.161 | -0.217 | -0.106 |
| | | stand_time | -0.008 | -0.029 | 0.013 |
| | | stressed1 | 0.068 | 0.045 | 0.09 |
| | | ts_length | -1.345 | -1.378 | -1.311 |
| | | search_effort | -0.013 | -0.038 | 0.013 |
| | | stand_time:stressed1 | -0.031 | -0.061 | -0.001 |
| | | stand_time:ts_length | -0.002 | -0.051 | 0.047 |
| | | stressed1:ts_length | -0.025 | -0.073 | 0.023 |
| | | stand_time:search_effort | -0.002 | -0.036 | 0.032 |
| | | stressed1:search_effort | 0.214 | 0.178 | 0.25 |
| | | ts_length:search_effort | 1.325 | 1.271 | 1.379 |
| | | stand_time:stressed1:ts_length | 0.543 | 0.473 | 0.614 |
| | | stand_time:stressed1:search_effort | -0.002 | -0.05 | 0.046 |
| | | stand_time:ts_length:search_effort | 0.01 | -0.069 | 0.089 |
| | | stressed1:ts_length:search_effort | -0.356 | -0.433 | -0.279 |
| | | stand_time:stressed1:ts_length:search_effort | -0.265 | -0.377 | -0.152 |
| | | *Random effects* | | | |
| | | Precision for comm_id | 54.154 | 53.56 | 54.734 |
| | | Precision for ar_id | 16.051 | 15.963 | 16.14 |
| | | Precision for time | 12.685 | 12.545 | 12.825 |

**Table S16.** Parameter estimates for the influence of community size, time series length, search effort and stress category on the probability of the multivariate Jacobian index exceeding its stability threshold of '1'. Median and upper and lower credible intervals are reported.

| Metric | Parameter | Median | CI_low | CI_high |
|---|---|---|---|---|
| multiJI | *Fixed effects* | | | |
| | (Intercept) | 1.138 | 0.847 | 1.429 |
| | n_spp | 0.112 | 0.093 | 0.13 |
| | stressed1 | 0.78 | 0.545 | 1.016 |
| | ts_length | -2.813 | -3.132 | -2.494 |
| | search_effort | 0.682 | 0.424 | 0.94 |
| | n_spp:stressed1 | -0.027 | -0.044 | -0.01 |
| | n_spp:ts_length | -0.146 | -0.168 | -0.124 |
| | stressed1:ts_length | -1.029 | -1.489 | -0.569 |
| | n_spp:search_effort | 0.005 | -0.014 | 0.024 |
| | stressed1:search_effort | 0.898 | 0.494 | 1.302 |
| | ts_length:search_effort | -0.316 | -0.833 | 0.201 |
| | n_spp:stressed1:ts_length | 0.055 | 0.023 | 0.087 |
| | n_spp:stressed1:search_effort | -0.027 | -0.056 | 0.003 |
| | n_spp:ts_length:search_effort | -0.051 | -0.087 | -0.015 |
| | stressed1:ts_length:search_effort | -0.26 | -1.027 | 0.507 |
| | n_spp:stressed1:ts_length:search_effort | 0.091 | 0.037 | 0.145 |
| | *Random effects* | | | |
| | Precision for comm_id | 3.723 | 2.628 | 5.105 |

**Table S17.** Parameter estimates for the influence of community size, time series length, search effort and stress category on the probability of the largest univariate Jacobian index exceeding its stability threshold of '1'. Median and upper and lower credible intervals are reported.

| Metric | Parameter | Median | CI_low | CI_high |
|---|---|---|---|---|
| max_uniJI | *Fixed effects* | | | |
| | (Intercept) | 0.541 | 0.331 | 1.429 |
| | n_spp | 0.034 | 0.021 | 0.13 |
| | stressed1 | 0.359 | 0.152 | 1.016 |
| | ts_length | 0.703 | 0.399 | -2.494 |
| | search_effort | 0.098 | -0.125 | 0.94 |
| | n_spp:stressed1 | -0.003 | -0.016 | -0.01 |
| | n_spp:ts_length | -0.031 | -0.05 | -0.124 |
| | stressed1:ts_length | 0.629 | 0.151 | -0.569 |
| | n_spp:search_effort | 0.023 | 0.008 | 0.024 |
| | stressed1:search_effort | 1.488 | 1.123 | 1.302 |
| | ts_length:search_effort | 1.414 | 0.894 | 0.201 |
| | n_spp:stressed1:ts_length | 0.034 | 0.001 | 0.087 |
| | n_spp:stressed1:search_effort | -0.033 | -0.059 | 0.003 |
| | n_spp:ts_length:search_effort | 0.021 | -0.014 | -0.015 |
| | stressed1:ts_length:search_effort | -2.613 | -3.462 | 0.507 |
| | n_spp:stressed1:ts_length:search_effort | 0.111 | 0.049 | 0.145 |
| | *Random effects* | | | |
| | Precision for comm_id | 8.559 | 5.99 | 5.105 |

**Table S18.** Parameter estimates for the influence of community size, time series length, search effort and stress category on the probability of multivariate autocorrelation exceeding its stability threshold of '0'. Median and upper and lower credible intervals are reported.

| Metric | Parameter | Median | CI_low | CI_high |
|---|---|---|---|---|
| max_uniJI | *Fixed effects* | | | |
| | (Intercept) | -0.722 | -1.376 | -0.064 |
| | n_spp | 0.155 | 0.117 | 0.193 |
| | stressed1 | 1.038 | 0.752 | 1.324 |
| | ts_length | -9.593 | -10.263 | -8.923 |
| | search_effort | 1.688 | 1.34 | 2.036 |
| | n_spp:stressed1 | -0.073 | -0.089 | -0.057 |
| | n_spp:ts_length | -0.19 | -0.226 | -0.153 |
| | stressed1:ts_length | 2.818 | 2.08 | 3.555 |
| | n_spp:search_effort | -0.106 | -0.125 | -0.087 |
| | stressed1:search_effort | 0.702 | 0.257 | 1.146 |
| | ts_length:search_effort | 3.56 | 2.691 | 4.43 |
| | n_spp:stressed1:ts_length | 0.138 | 0.097 | 0.179 |
| | n_spp:stressed1:search_effort | 0.055 | 0.03 | 0.081 |
| | n_spp:ts_length:search_effort | 0.269 | 0.221 | 0.316 |
| | stressed1:ts_length:search_effort | -0.689 | -1.699 | 0.321 |
| | n_spp:stressed1:ts_length:search_effort | -0.184 | -0.241 | -0.126 |
| | *Random effects* | | | |
| | Precision for comm_id | 0.595 | 0.417 | 0.82 |
